# SLiM 5: Eco-evolutionary simulations across multiple chromosomes and full genomes

**DOI:** 10.1101/2025.08.07.669155

**Authors:** Benjamin C. Haller, Peter L. Ralph, Philipp W. Messer

## Abstract

Evolutionary simulations of multiple chromosomes, even up to the scale of full-genome simulations, are becoming increasingly important in population genetics and evolutionary ecology. Unfortunately, the popular simulation framework SLiM has always been intrinsically limited to simulations of a single diploid chromosome. Modeling multiple chromosomes of different types, such as sex chromosomes, has always been cumbersome even with scripting, presenting a substantial barrier to the development of full-genome simulations. Here we present SLiM 5, a major extension of SLiM’s capabilities for simulating multiple chromosomes. Modeling up to 256 chromosomes is now possible, and each chromosome may belong to any of a wide variety of types – not just autosomes (diploid and haploid), but also sex chromosomes (X, Y, Z, and W), haploid mitochondrial and chloroplast DNA, and more. This new functionality is integrated across all of SLiM, including not only the mechanics of reproduction and inheritance, but also input and output of multi-chromosome data in formats like VCF, and tree-sequence recording across multiple chromosomes. New recipes in the SLiM manual demonstrate these new features, and SLiM’s graphical modeling environment, SLiMgui, has been extended in many ways for the visualization of multi-chromosome models. These new features will open new horizons and enable a heightened level of realism for full-genome simulations.

## Introduction

Evolutionary simulations are an increasingly central tool for fields such as population genetics, evolutionary ecology, and conservation biology. Within that general trend comes a demand for the ability to simulate multiple chromosomes and even full genomes, driven by several factors. One is the increasing availability of full-genome sequence data for individuals, across a variety of study systems; with such data at hand, it may often be desirable to plug it into a simulation, or simulate at a scale that matches the data (Carvajal-Rodriguez, 2010; Li & Durbin, 2011; Hoban et al. 2012). Another is that analyses of empirical data are being done at full-genome scale, such as GWAS (Tam et al., 2019; Uffelmann et al., 2021) and selection scans (Oleksyk, Smith, & O’Brien, 2010; Haasl & Payseur, 2016), and so simulation at that scale is desirable to provide data for developing and testing these methods. In particular, simulation is increasingly an important tool for understanding complex, polygenic traits (Meuwissen & Goddard, 2010; Boyle, Li, & Pritchard, 2017; Evans et al., 2018), for which causal loci are often spread out across all chromosomes; simulations of single chromosomes must therefore either understate heritability or overstate linkage between causal loci. Furthermore, simulating multiple chromosomes or even full genomes can be important because loci on different chromosomes can interact, such as in mitonuclear interactions (Baris et al., 2017; Zaida & Makova, 2019; Munasinghe, Haller & Clark, 2022); because linkage disequilibrium can exist between loci on different chromosomes even though they are physically unlinked (Oraguzie et al., 2007; Rohlfs, Swanson, & Weir, 2010; Skelley, Magwene, & Stone, 2016); and because the different patterns of inheritance exhibited by autosomes, sex chromosomes, and other types of DNA (e.g., mitochondrial, chloroplast) can be important to evolutionary dynamics (Birky, 2001; Korpelainen, 2004; Mank, 2012; Wilson Sayres, 2018; Vicoso, 2019). Finally, understanding the genetic and evolutionary basis of sex differences (in, for instance, health outcomes) requires jointly considering autosomes and sex chromosomes (Gilks, Abbott, & Morrow, 2014; Khramtsova, Davis, & Stranger, 2019; Wilson & Buetow, 2020; Massey et al., 2021).

SLiM is a popular and widely used eco-evolutionary simulation framework that we have continuously maintained, supported, and improved for more than a decade (Messer, 2013; Haller & Messer, 2017; Haller & Messer, 2019; Haller & Messer, 2023). Because it is scriptable, using a language called Eidos, it is quite flexible, and can model a wide variety of ecological and evolutionary dynamics. However, SLiM has never had proper built-in support for simulating multiple chromosomes. One could model unlinked loci, using a recombination rate map with a rate of 0.5 between one base position and the next such that the simulated chromosome was divided into two (or more) unlinked portions, but this had many limitations. For example, it was not possible to make one of those “effective chromosomes” an autosome, another an X, and another a Y without complex, technical, and error-prone scripting, and even then, functionality such as base positions within chromosomes, gene conversion tracts at the edges of chromosomes, VCF input and output involving the chromosomes, and visualization of multiple chromosomes in SLiMgui would not be handled correctly. Full-genome simulations with sex chromosomes (and perhaps other elements such as mitochondrial DNA) were therefore not quite impossible, but were unsupported and prohibitively difficult in practice, and this has been limiting for many potential applications of SLiM.

Here, we introduce SLiM 5, which fundamentally extends SLiM’s capabilities in this area. With SLiM 5, up to 256 chromosomes (per species) can be simulated explicitly. Each can be of a different type, chosen from among a broad set of supported types – autosomes (diploid and haploid), sex chromosomes (X, Y, Z, and W), and a variety of haploid chromosome types with special inheritance patterns in sexual models (e.g., present in both sexes or only one, inherited from one sex or from the other) that can be used to model things like mitochondrial and chloroplast DNA; a full list is provided in Table 1. SLiM handles all of the complexities of ploidy, reproduction, and inheritance internally; the user simply has to declare the chromosomes they want to implement, and supply information such as their lengths, types, and recombination and mutation rates. With this architecture it is simple, for example, to construct a full-genome model of dragonflies with an X0 system (males X0, females XX, where the 0 denotes the lack of a Y chromosome), or of cichlid fish in the genus *Tilapia* (males XYZZ, females XXZW). This new support for multiple chromosomes extends across all of SLiM, from VCF input and output to graphical model visualizations in SLiMgui. Even tree-sequence recording (Kelleher et al., 2018; Haller et al., 2019), which can record the pattern of ancestry along the chromosome in SliM simulations, now works with multi-chromosome models; SLiM will provide a separate recorded tree sequence for each chromosome in the model, saved as separate `.trees` files in a new multi-chromosome format called a *trees archive*. In short, SLiM 5 makes it easy to build multi-chromosome models, up to the scale of full genomes.

**Table 1.**
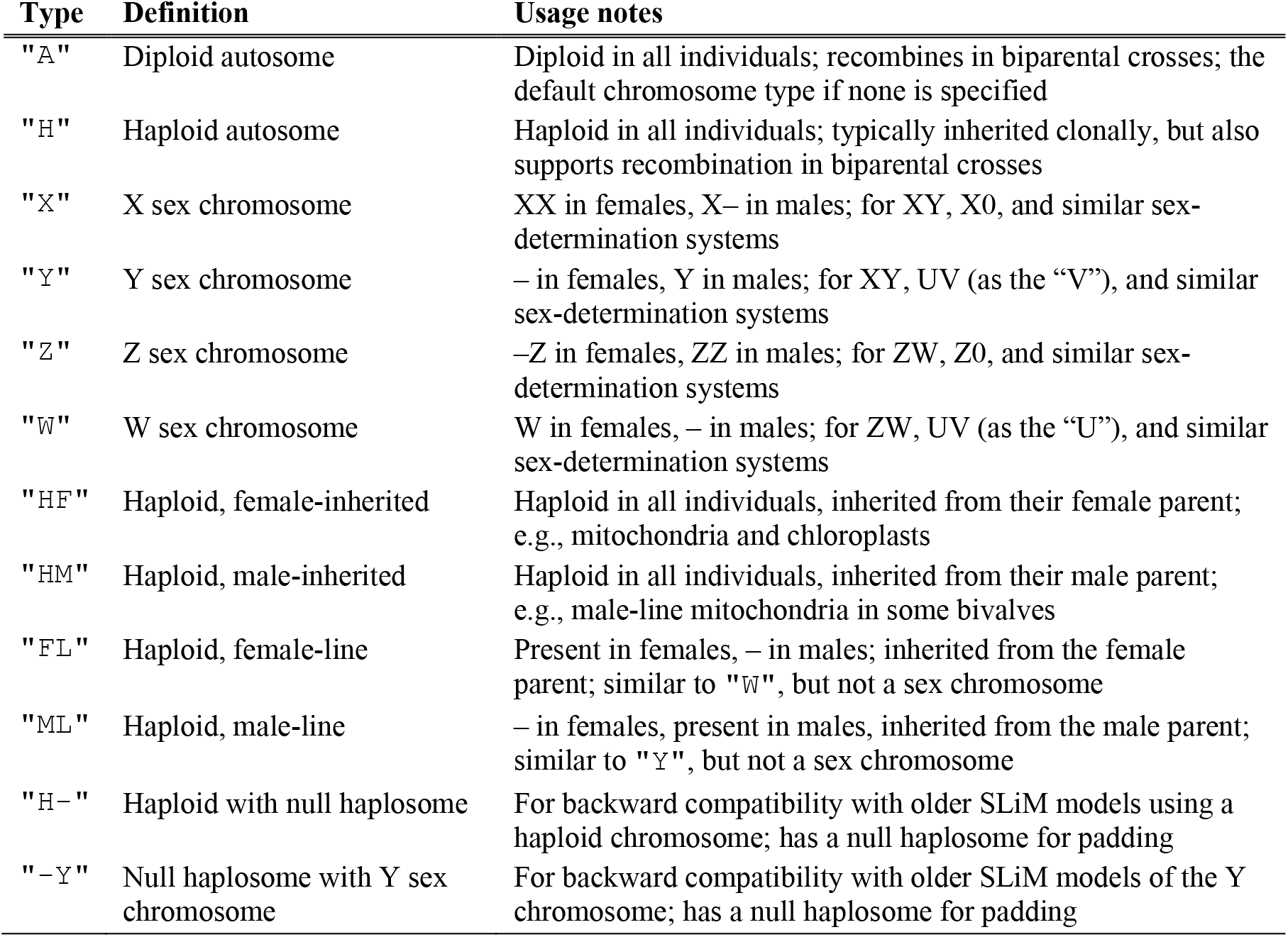
Chromosome types supported in SLiM 5, with notes on their usage. A dash, –, represents a null haplosome that is used as a placeholder.

### On Haplosomes

Before we get into describing SLiM’s new multi-chromosome features in more detail, there is an essential point of terminology that must be addressed. In previous versions of SLiM the genetic model was fairly simple. Each diploid individual would possess two copies of the simulated chromosome; each of those copies was called a “genome”, so a diploid individual would contain two “genomes”, which would contain the genetic information for that individual – which mutations it possessed, in particular (Fig. 1A).

**Figure 1.**
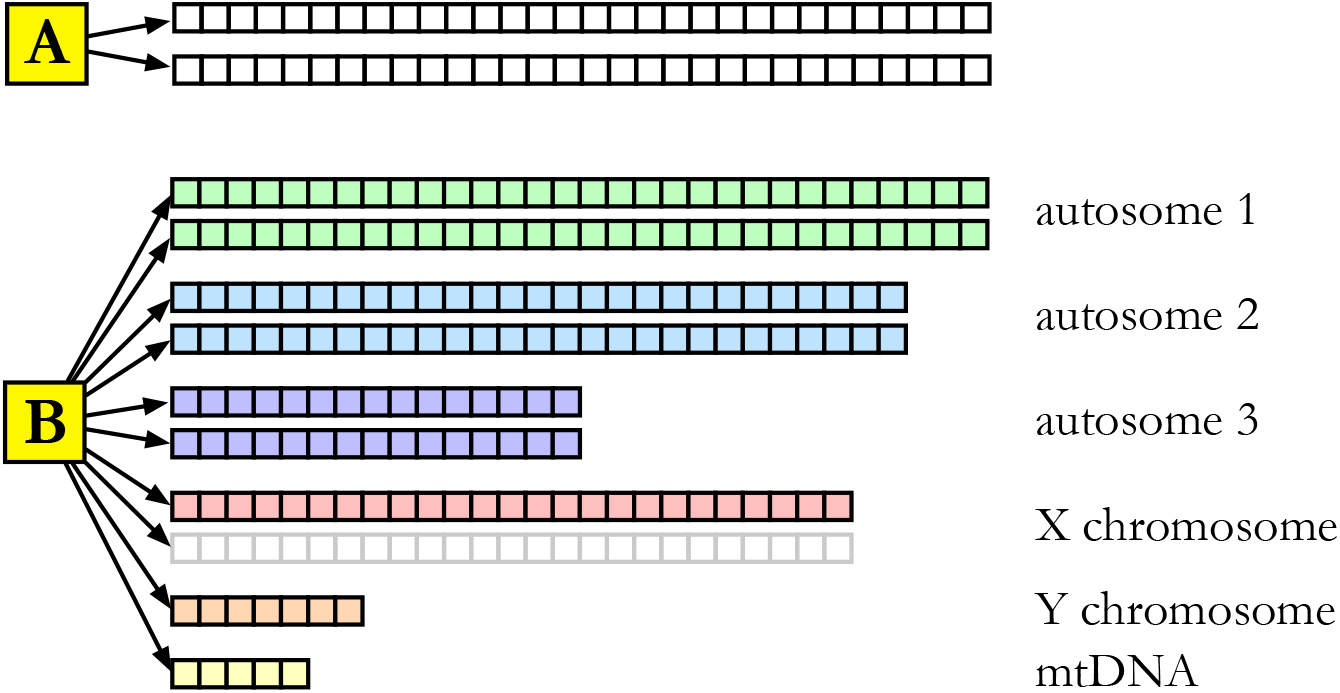
(A) Schematic of the genetic structure in SLiM 4 and earlier. One individual is represented by the large yellow square, at left, labeled **A**. That individual contains two “genomes” represented as strips of boxes connected to the individual by arrows; each box represents one base position. Each box – each base position within a genome – could contain a mutation in a given individual. Each individual thus contains two “genomes” that together hold all of that individual’s genetic information. (B) Schematic of the top-level genetic structure in SLiM 5. One individual, labeled **B**, can now contain many of the things that used to be called “genomes”, now called “haplosomes”; different haplosomes are associated with different chromosomes, potentially occurring at different ploidies in different individuals (perhaps depending upon the individual’s sex, as here). The second copy of the X chromosome, for example, is shown in gray because it is not used in this male individual; in SLiM this is represented by a placeholder object called a null haplosome.

SLiM 5 extends this conceptual model to multiple chromosomes. Individuals can now contain genetic information associated with many different chromosomes, in varying numbers of haploid copies of each chromosome, depending upon ploidy (Fig. 1B). The word “genome” clearly can no longer be used to refer to these individual haploid copies. Remarkably, there does not actually seem to be a short-hand term in biology for a “haploid copy of a given chromosome”. Instead, one usually just says, for example, that “females have two homologous copies [or versions] of the X chromosome”, or one might refer to “the maternal copy [or version] of chromosome 3” in an individual. However, to keep the names of objects, functions, methods, and properties manageable and precise in SLiM, we felt the need to declare a distinct and concise term that refers to a “single copy of a given chromosome”. After considering various alternatives, we ultimately settled upon “haplosome”. A “haplosome”, then, is defined as “one of the homologous versions of a given chromosome in a given individual”. The Greek root for the first half, “haplóos” (ᾰ̔πλóΟς), means “single” (as in “haploid”); the second half, “sôma” (σῶμα), means “body” (as in “chromosome”). A “haplosome” is thus a “single body” that carries one genetic sequence in an individual. For a diploid chromosome, an individual has two haplosomes; for a haploid chromosome, it has one haplosome. The term has meaningful echoes of other terms in biology, fits the concept well, and is, rather serendipitously, available.

We acknowledge that coining a new term for such a central concept is no small thing, and could receive pushback from some users. However, our rationale for introducing this terminology is that the previous usage of “genome” was clearly no longer workable, and that referring to a “copy of a chromosome” would have been too verbose and also too confusing, given that this term would be used all across SLiM (as the name of a class, formerly the `Genome` class and now the `Haplosome` class, and in property names, method names, and so forth). We thus hope that the community will, in the end, embrace the term “haplosome”.

### A Simple Example

For a simple first example of a multi-chromosome model in SLiM 5, let’s look at a simple hermaphroditic model with two diploid autosomes, just to show the basic approach and introduce some SLiM concepts. The script for this model is shown in Code Sample 1.

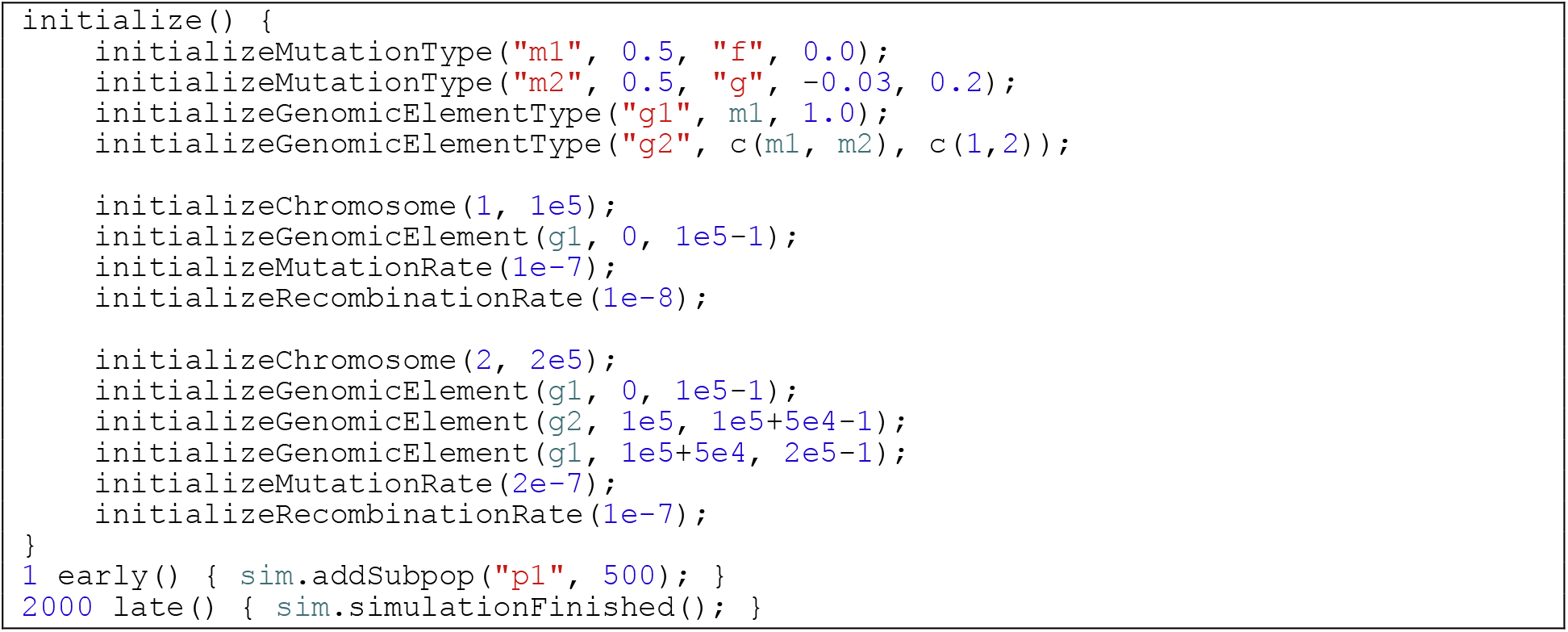

Code Sample 1. This is the file `SimpleExample.slim` in our repository.

Most of the script is in the form of an `initialize()` callback, a block of code that initializes the basic structure of the simulation. Here, that initialization is about setting up the model’s genetic structure. It begins by defining two “mutation types”, with calls to `initializeMutationType()`. A mutation type is essentially a distribution of fitness effects (DFE). The first, `m1`, is configured to have a fixed (`“f”`) selection coefficient of `0.0`; it has a dominance coefficient of `0.5`, but given that it is neutral, that doesn’t matter. So `m1` represents neutral mutations. The second, `m2`, draws selection coefficients from a gamma distribution with a mean of `-0.03` and a shape parameter of `0.2`. (Gamma distributions cannot have a negative mean, but this tells SLiM to use a mean of 0.03 and then flip the sign of all draws.) So `m2` represents a particular class of deleterious mutations.

Next it defines two “genomic element types”, categories of genomic regions. The first, `g1`, draws mutations only from the `m1` DFE, representing neutral non-coding regions. The second, `g2`, draws mutations from `m1` and `m2` in a 1:2 ratio, representing coding regions in which, as an approximation, one-third of mutations are neutral and two-thirds are deleterious.

And finally, the `initialize()` callback defines the two chromosomes, using the genomic element types `g1` and `g2`. A call to `initializeChromosome()` defines each chromosome; the first parameter is an integer identifier for the chromosome (`1` and `2`, here), and the second is a length in base positions (`1e5` and `2e5`). After each call to `initializeChromosome()`, SLiM expects further calls that configure that chromosome. Here, calls to `initializeGenomicElement()` set up regions of the chromosome that are non-coding (using `g1`) or coding (using `g2`), and then a mutation rate and recombination rate is set up for the chromosome. Calls to `initializeGenomicElement()`, `initializeMutationRate()`, and `initializeRecombinationRate()` affect the most recently initialized chromosome, rather than explicitly taking a `chromosome` argument.

That configuration is most of the model. There is then an event that runs early in the first tick (the first timestep) of the model’s execution (`1 early()`), which creates a new subpopulation named `“p1”` with `500` individuals, and an event that runs late in tick 2000 (`2000 late()`), which explicitly ends the simulation with a call to `sim.simulationFinished()`. Note that `sim` is a built-in object in SLiM that represents the simulation (or more precisely, the single species being simulated, in a single-species model like this). Basically, `sim` is a top-level object that is used to control the simulation of the species, and here we call methods of it to add a subpopulation, and then 2000 ticks later, to end the simulation. Ticks are the basic time unit of SLiM; in a Wright–Fisher model such as this, in each tick the parental generation reproduces to generate a set of offspring, and then the parents die and the offspring become the new parents for the next tick, so one tick equals one generation for the species being simulated.

Almost all of this is standard SLiM code that would have run in SLiM 2, 3, or 4; the twist here is the use of `initializeChromosome()` to define two separate autosomes, with different lengths, different configurations of coding and non-coding regions, and different mutation and recombination rates. If we run this model in SLiMgui, SLiM’s graphical modeling environment, the chromosome view in the simulation window shows the genetic structure of the species (Fig. 2).

**Figure 2.**
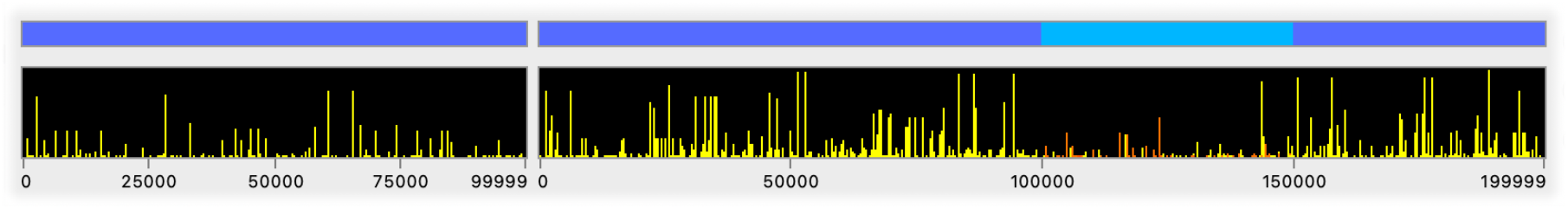
The chromosome view’s display for the simple model in Code Sample 1, after executing for 2000 ticks. Both chromosomes are displayed here. The top strip of the display is the “overview”, showing the basic genetic structure of the model. Here, the first chromosome has a single genomic element of type `g1`, defining a non-coding region that undergoes only neutral mutations, whereas the second chromosome has three genomic elements, of types `g1`, `g2`, and `g1`, defining a coding region flanked by non-coding regions. The taller strip below is the “detail” view, which shows the mutations present in the population. Each vertical bar in the detail view represents one mutation, and the height of the bar represents the mutation’s frequency in the population; tall bars are high-frequency mutations, short bars are low-frequency. The yellow bars represent neutral mutations; yellow is the color of neutrality in SLiM. In the first chromosome, which is entirely neutral, all mutations are yellow; in the second chromosome, some deleterious mutations (shown in orange and red) are present within the central coding region.

With this simple setup, we already have a two-chromosome model with some genetic structure, both neutral and deleterious mutations, and (although it can’t be seen in Fig. 2) different mutation and recombination rates for the two chromosomes. This could be extended to have more mutation types representing more DFEs (beneficial mutations, for example), more genomic element types representing different kinds of genomic regions (introns versus exons versus regulatory regions, for example), and more chromosomes with a more complex genetic structure. We could also have multiple subpopulations, or we could make this a continuous-space model with local interactions between individuals; and we could have SLiM model explicit nucleotides, and load an ancestral sequence for each chromosome from a FASTA file and save out a VCF file at the end containing the segregating SNPs; and so forth. In other words, all of the features of previous versions of SLiM remain available; but in addition to that, we now have two chromosomes, which we can configure separately, and which SLiM manages for us.

### A Full-Genome *Drosophila* Model

SLiM now supports many different kinds of chromosomes beyond the autosomes used in the previous section (Table 1). To demonstrate this, we now construct a whole-genome model of *Drosophila melanogaster*, including its sex chromosomes and mitochondrial DNA.

Most of the model is shown in Code Sample 2 (the rest will be shown in the next section). In some ways it is quite similar to what we have seen before. Again, almost all of the model is the configuration of the genetics, since that is what we are focusing on. At the beginning, this model uses the `readCSV()` function to load recombination rate maps from files on disk. This function returns a `DataFrame` object, much like a data frame in R, containing columns of data. In this case, each `DataFrame` contains a column of recombination rates, and a column of base positions at which each rate ends. These rate maps are based on empirical recombination rate data from Comeron et al. (2012); they have only been modified slightly, primarily to tack the 2L and 2R arms together, and the 3L and 3R arms together, since we will treat those chromosomes as a whole. These files, and the procedure by which they were generated, are provided in our repository in the `recmaps` directory.

Next, we get the length of each chromosome (including the mtDNA, which we will call a “chromosome” henceforth, following SLiM’s usage). For the chromosomes that have recombination rate maps, we use the end position of the rate map as the chromosome length. Additional genome data are input here as obtained from `stdpopsim` (version 0.3.0; Adrion et al. 2020, Lauterbur et al. 2023): lengths of nonrecombining chromosomes (dos Santos et al., 2015; Hoskins et al., 2015) and mutation rates (Schrider et al., 2013). (Note that `stdpopsim` is itself a high-level simulator, which uses both `msprime` and SLiM as back ends, but it does not presently support multiple chromosomes.)

Next we call `initializeSex()` to let SLiM know this is a model with separate sexes; this will allow us to define sex chromosomes, and to set recombination rates that differ between females and males (as they do in *D. melanogaster*). We set up only a neutral mutation type, `m1`, and a genomic element type, `g1`, that represents neutral regions using `m1`. It would be straightforward to get the coding sequence annotation and a distribution of fitness effects for *D. melanogaster* from `stdpopsim`; see Haller, Nelson, & Messer (2025) for an example of that.

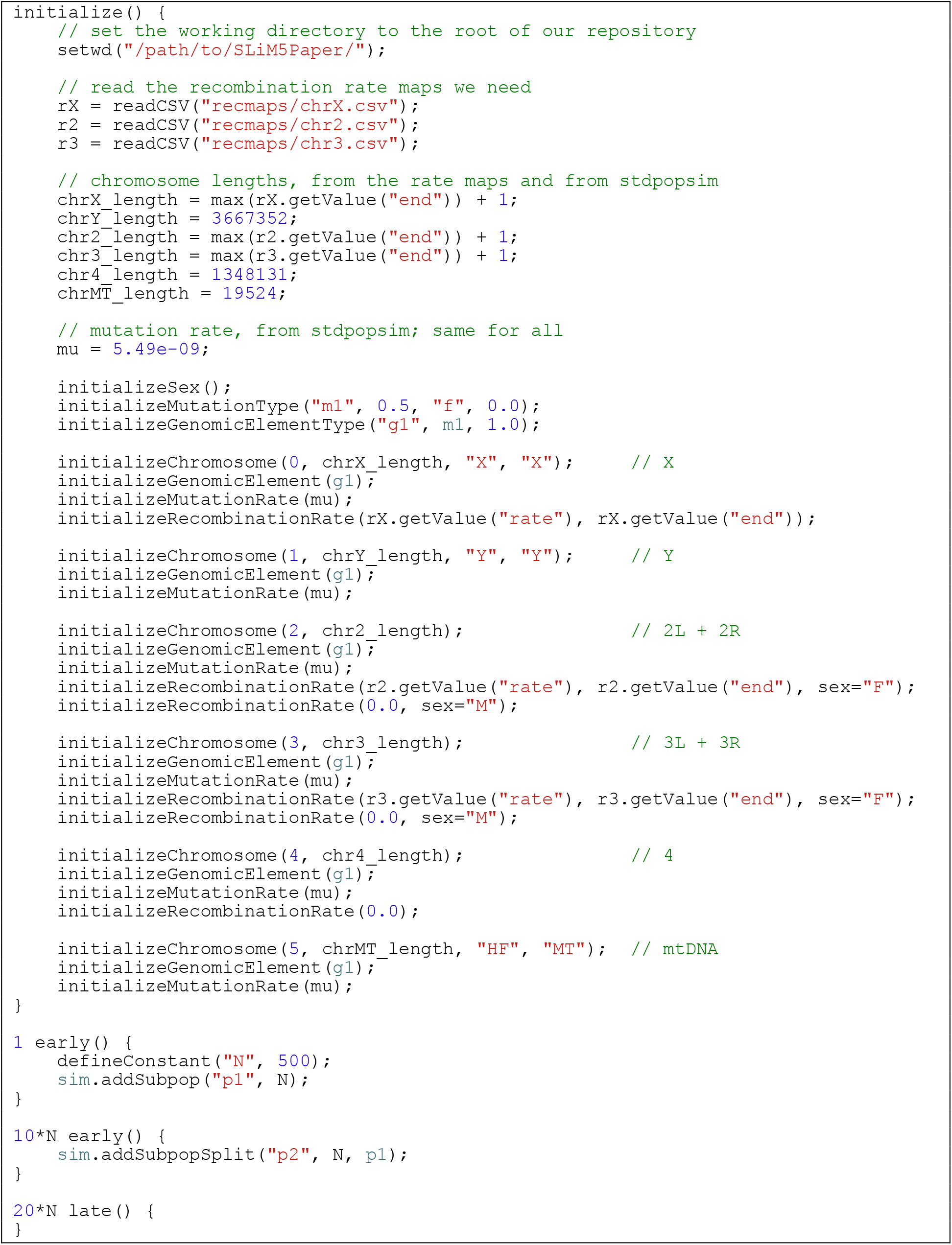

Code Sample 2. This is (most of) the file `DroMelFullGenome.slim` in our repository.

The remainder of the `initialize()` callback is very similar to what we did in Code Sample 1, with a few differences. First of all, in some of the `initializeChromosome()` calls we pass additional parameters beyond the `id` and `length` we saw before. The third parameter is `type`; we use it to specify the X, Y, and mtDNA chromosomes with `“X”`, `“Y”`, and `“HF”` (`“HF”` stands for “haploid female”, and means that it is a haploid chromosome that all individuals carry, inherited from the female parent). The fourth parameter is `symbol`; we use it to define a symbol, of type `string`, that is used by SLiM to refer to the chromosome, as we’ll see later. For the mtDNA, we use `“MT”` as the symbol.

A second difference is that we now allow `initializeGenomicElement()` to infer the span of the genomic element for us, by omitting the start and end positions. In that case, it assumes that the genomic element defined spans the entire chromosome, which is what we want since we are not setting up separate genomic elements for exons, introns, etc. This shorthand is just a minor convenience.

The third difference is how we specify the recombination rates. Before, we just used a simple call to `initializeRecombinationRate()` with a single rate like `1e-7` or `1e-8` (per base pair and meiosis). Here, we instead use the rate maps that we loaded at the start, originally from Comeron et al. (2012) – but only in the females, which we tell SLiM by passing `sex=“F”`. (Note that Comeron et al. (2012) provide their recombination rate maps based upon an experimentally observed number of crossovers per female meiosis, not as sex-averaged rates; since we then use the recombination rate maps only in females, no correction is needed.) We get the rates and end positions for each rate map out of the loaded `DataFrame` objects using the `getValue()` method (`“rate”` and `“end”` are the names of the columns in the loaded CSV files). In the males, we set a recombination rate of `0.0`, passing `sex=“M”`.

The `1 early()` event here sets up a single subpopulation in tick 1, `p1`. It is given a size `N`, where `N` is a constant that we define with `defineConstant()`. By making `N` a defined constant, we can use it elsewhere in the model as well.

The next event shows off another feature of SLiM 5 that is new since SLiM 4.0. We define this event as running in tick `10*N`, using the constant `N` that we defined before. Notice that `N` is not even defined at the end of the model’s `initialize()` callback, but SLiM doesn’t mind; it just waits until `N` becomes defined, and then schedules the event for us. This new feature is called “dynamic scheduling”.

By scheduling this event in tick `10*N`, we give the population time to (more or less) equilibrate. This provides a “burn-in period”, during which genetic diversity can build up until it reaches a state close to mutation–drift balance. Now, in tick `10*N`, we split off a new subpopulation using the `addSubpopSplit()` method to create subpopulation `p2`, also with size `N`, representing a vicariance event that separates the two subpopulations with no gene flow between them. The model ends with the `20*N late()` event; so it runs from `10*N` to `20*N` as two separate subpopulations.

If we run this model in SLiMgui, we can again look at the chromosome view to see what happened (Fig. 3). However, we would like to understand more about what is happening in this model over time. To this end, we will look at a very powerful visualization technique in the next section.

**Figure 3.**
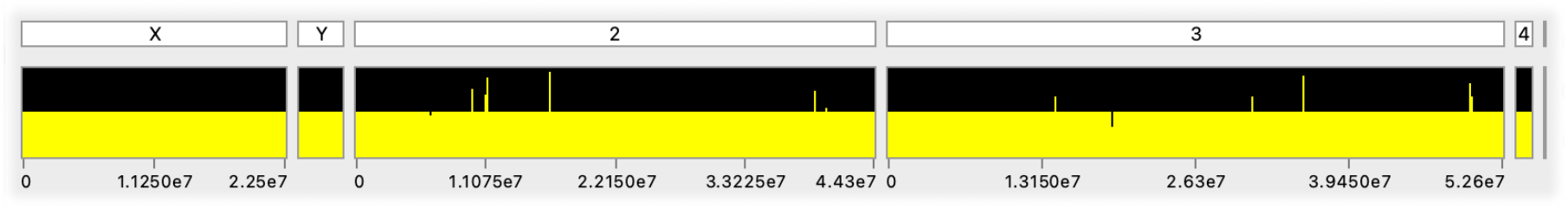
The chromosome view’s display for the *Drosophila melanogaster* model in Code Sample 2, after executing for 10*N* ticks following a 10*N* burn-in. Much of the genetic diversity is in the form of fixed differences between the two subpopulations in the model. Unlike in Fig. 2, the chromosome view is showing the names of the chromosomes (their `symbol` property values); the chromosome view switches to this alternate display mode when the mouse cursor briefly hovers over the chromosome overview. The mtDNA chromosome is the tiny sliver at the far right; we could zoom in on it (or any chromosome) in SLiMgui to see more detail, or even open a separate chromosome display window for it.

### Plotting *F*ST

The two subpopulations in this model are isolated throughout the vicariance period, and therefore build up a high level of neutral genetic differentiation between each other. The level of differentiation between populations is commonly measured with the *F*_ST_ statistic (Holsinger & Weir, 2009), and SLiM provides a built-in function, `calcFST()`, that can calculate it for us. We can extend our *D. melanogaster* model with Code Sample 3 to take advantage of this functionality.

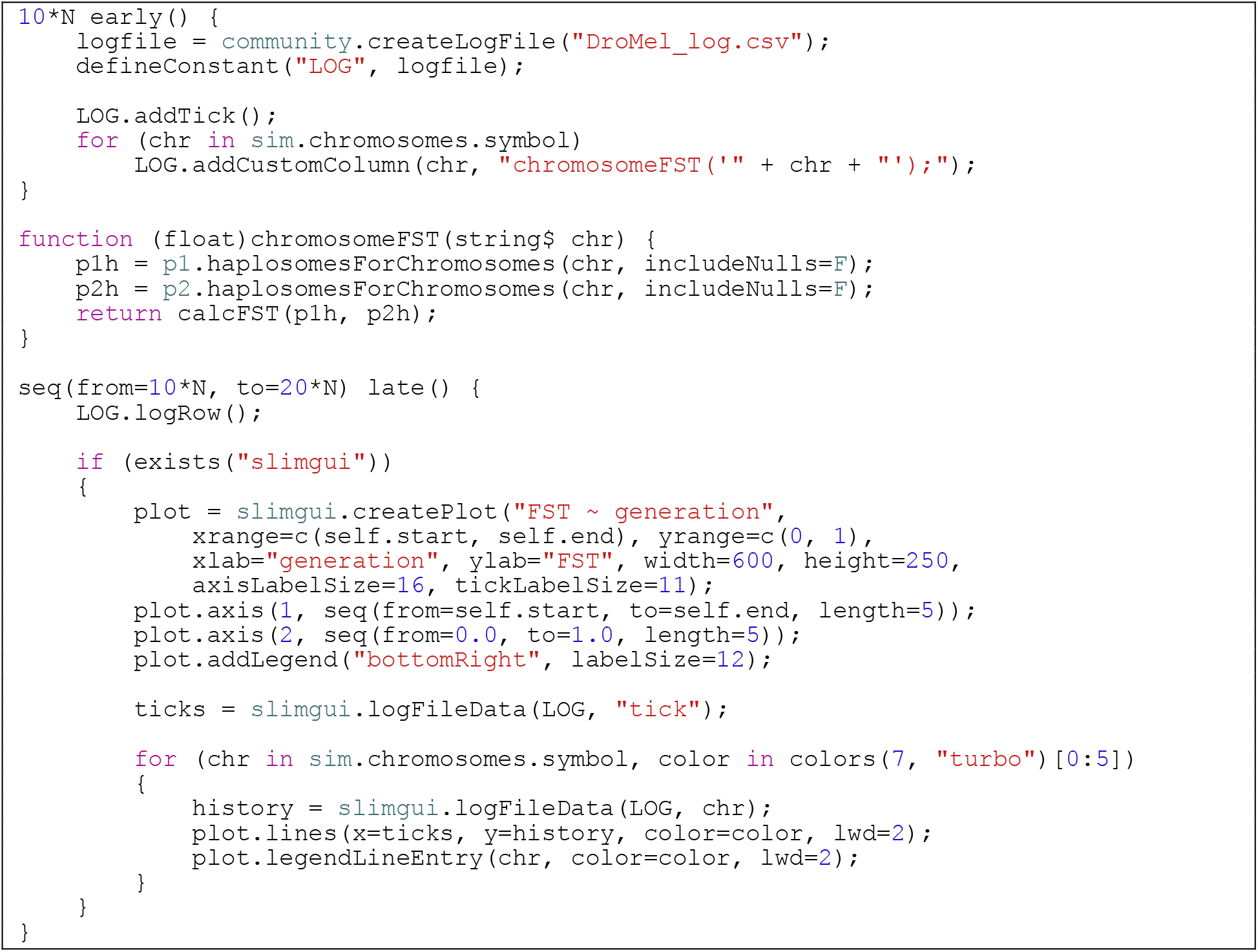

Code Sample 3. The remaining portion of `DroMelFullGenome.slim`.

The first event, `10*N early()`, sets up a `LogFile` object at the point when vicariance begins. `LogFile` is a class provided by SLiM that can log out information about your simulation to a file. The call to `createLogFile()` makes a new `LogFile` object, which we then put into a global constant, `LOG`, with `defineConstant()` so we can easily refer to it later on. The rest of this event configures what information will be logged; the output of a `LogFile` is configured just once, at the time it is created, and then it can log out a new row of data, following that configuration, either automatically (with some specified period) or on demand. Here we configure it so that each time it logs a new row, it logs out the current tick (at the time that the row is logged) as the first column of data. Then the `for` loop configures the `LogFile`, with calls to `addCustomColumn()`, so that it logs out custom data for each of the chromosomes in the model. The `addCustomColumn()` method allows the user to provide a little snippet of Eidos code called a “lambda”, given as a string; here, the lambda calls a function named `chromosomeFST()`, passing it the symbol for the chromosome.

The `chromosomeFST()` function is not built in; in the next script block, we actually define it as a new user-defined function. The function declaration syntax here states that `chromosomeFST()` is a new function that takes a single string as a parameter (assigned the name `chr`), and returns a `float` – a floating-point number, a non-integer number. The body of this function is quite simple; it calls the `haplosomesForChromosomes()` method on `p1` and `p2` to find all of the haplosomes associated with the chromosome that was passed in to the function. (Passing `includeNulls=F` tells the method to exclude the “null haplosomes” that act as placeholders for the second X haplosome in males, and for the Y haplosome in females; we don’t want to include those in the *F*_ST_ calculations, since an error would result.) In the last line of the function, we call `calcFST()` with those two sets of haplosomes; SLiM does the *F*_ST_ calculation for us, and we return the result. That result will get returned back to our `LogFile` object, since this function gets called to generate the data logged out for the columns that we added with `addCustomColumn()`. The end result, then, is that the `LogFile` will log out *F*_ST_ values for each of the chromosomes each time that it logs a new row of data.

The last event in Code Sample 3 code uses dynamic scheduling again. Dynamic scheduling can involve calls to functions, and indeed, the schedule for this event is generated by the `seq()` function, which generates a sequence of numbers from a starting value (`10*N`) to a final value (`20*N`). The event will therefore run from tick `10*N` to `20*N`, plotting data for every generation during the vicariance period.

The first line of this event calls the `logRow()` method of our `LogFile` object, which tells it to immediately log out a new row of data to its file. (Behind the scenes, this will trigger a call to `chromosomeFST()` for each chromosome, following the configuration we set up before.) So a new row of data gets logged each tick, after the burn-in period has ended.

After that, this event generates a custom plot which will update in SLiMgui in every tick after the burn-in period ends, showing *F*_ST_ over time for each chromosome. Let’s look at the final plot, at the end of a run of this model (Fig. 4), before we discuss the code that generates the plot.

**Figure 4.**
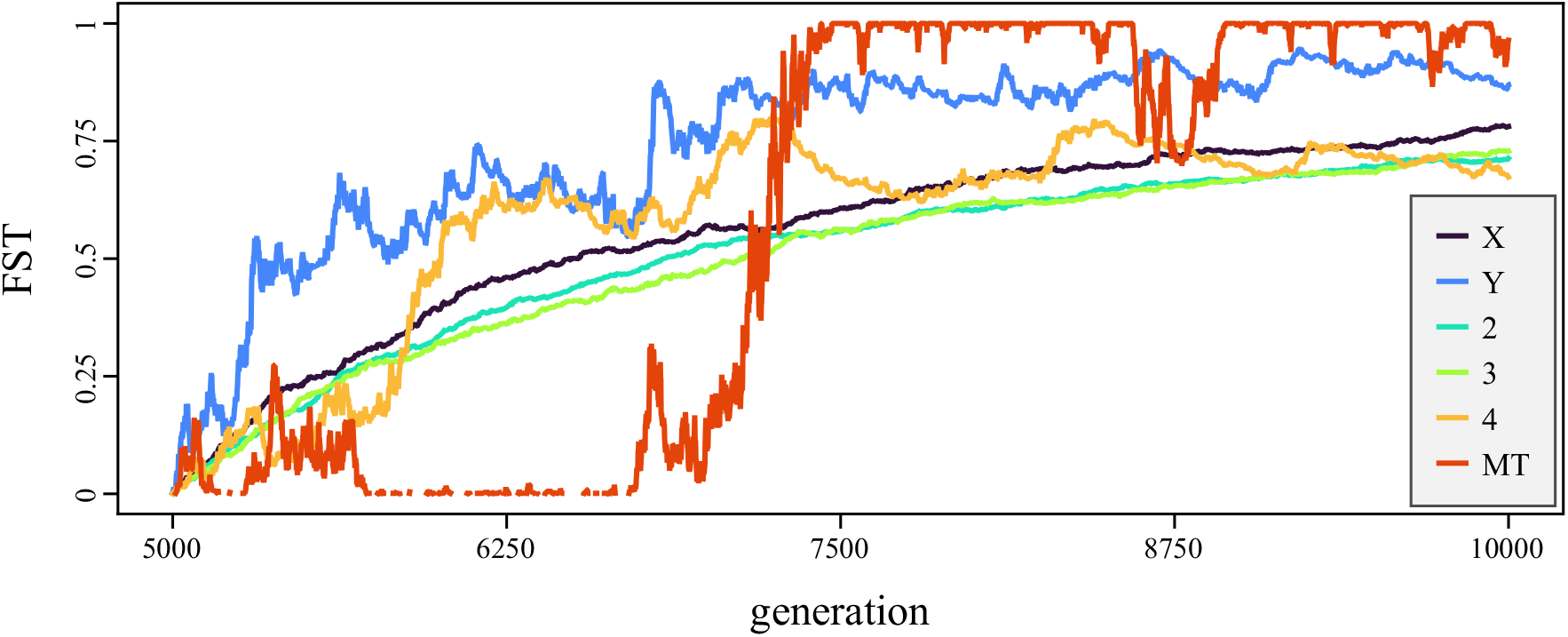
This plot shows *F*_ST_ as a function of time (generation), for each of the chromosomes in our *D. melanogaster* model, beginning in tick 5000 (the end of burn-in) until tick 10000 (the end of the simulation). In general, *F*_ST_ levels start out very low (because the population began as a single population that split in two) and then rise towards an equilibrium value of 1.0. The large chromosomes (X, 2, and 3) show the smoothest increases; chromosome 4, which is very short and does not recombine, has a somewhat bumpier trajectory but largely equilibrates in the same way as X, 2, and 3. The Y and MT chromosomes, which are haploid and have a smaller effective population size (*N*_e_), have much bumpier trajectories. Note that this plot was generated directly as a PDF by SLiMgui from the script in Code Sample 3; it has not been modified or retouched.

Custom plotting is another feature that is new since SLiM 4.0, and it can be very useful for model exploration and visualization. Sometimes, as here, it can even generate polished plots that are ready for publication. The plotting code runs only under SLiMgui, because drawing plots uses the Qt widget kit that SLiMgui uses for its user interface; the `slim` command-line tool does not link against Qt, so plotting is unavailable there. (`LogFile` will still log out *F*_ST_ values to disk, though.) The presence of SLiMgui is checked with the `exists()` function, taking advantage of the fact that a global object named `slimgui` exists only when running under SLiMgui. In that case, we make a new plot by calling the `createPlot()` method on that `slimgui` object, requesting various fairly self-explanatory options. Note that we create a new plot every tick, so that the plot updates as the model is running; unless the plot is quite complicated, this is very fast, and barely slows the SLiMgui down. For a more aesthetic display, we then configure the axis tick positions, and add a legend to the plot.

This plotting code uses the data collected by our `LogFile`; when running under SLiMgui, all of that data is kept in memory and can be accessed with the `logFileData()` method of the `slimgui` object. (When running at the command line, logged data is not kept in memory, to reduce the memory footprint of the `slim` process.) The initial call to `logFileData()` gets the ticks in which logging has been done so far; the column named `“tick”` is produced as a result of the `addTick()` configuration call we made during the setup of the `LogFile`.

The colored lines in the plot are produced by the `for` loop that follows. Note that this `for` loop iterates in synchrony over two vectors: the chromosome symbols, using loop index variable `chr`, and a vector of plotting colors, using loop index variable `color`. Each chromosome thus has an associated color, within the body of the `for` loop. This is called a “joint `for` loop”, and is another feature added since SLiM 4.0 – an extremely useful and convenient feature, for situations like this!

The loop starts by fetching the *F*_ST_ history for the current chromosome in the loop, with a call to `logFileData()`. Then it adds a line to the plot showing the *F*_ST_ history for that focal chromosome, and adds an entry in the plot legend with the correct label and color.

Note that custom plots like this can incorporate plotted points as well as lines, and can even write text into the plot window – for example, to add labels in particular spots. Custom plotting in SLiMgui is flexible enough for many purposes, although for particularly complex plots one might still prefer R, Python, or other packages with more sophisticated plotting capabilities.

## Discussion

SLiM 5 introduces support for simulating multiple chromosomes, up to the scale of full genomes, including sex chromosomes and genetic entities such as mitochondrial and chloroplast DNA. It handles all of the details of inheritance for different chromosome types; once a set of chromosomes has been defined, SLiM can manage the rest of the simulation automatically unless there is a need for specific control over recombination behavior, different reproductive modes, or other such complexity (which can be handled in script). Furthermore, its graphical modeling environment, SLiMgui, has been updated to display multiple chromosomes, and tree-sequence recording can separately record the ancestry along each chromosome; support for multiple chromosomes has been deeply integrated across all of SLiM. We illustrated these new capabilities with two example models that demonstrate how to construct multi-chromosome and full-genome models in SLiM 5.

Limitations remain in SLiM’s genetic model, however. One major limitation is that ploidies above diploidy, such as tetraploidy, are not yet supported. This is a difficult area since many aspects of a tetraploid simulation, from recombination to fitness calculations, would differ substantially from SLiM’s current paradigm. Nevertheless, SLiM 5’s new design does pave the way for possible future support for higher ploidies, which could perhaps now fit into SLiM more easily as a new chromosome type; at present, all of the chromosome types supported by SLiM are either haploid or diploid, but one can imagine extending that design to add a new tetraploid chromosome type, with four haplosomes per individual, as long as other aspects of SLiM were also extended to support that configuration.

Another remaining limitation regards chromosomes that can exist in a wide variety of ploidies in different individuals. These are often called “B-chromosomes”, “supernumerary chromosomes”, or “accessory chromosomes”, and occur in many species (Camacho, Sharbel, & Beukeboom, 2000). It is hard to imagine SLiM ever supporting these intrinsically, given how variable they are; but since SLiM is scriptable, one could already script simple things like tracking the number and type of B-chromosomes present in each individual and associating a fitness effect with each type. Explicitly tracking the genetics of each B-chromosome copy in each individual seems difficult to fit into SLiM’s model, however.

A third limitation is that at present, SLiM 5 does not have intrinsic support for modeling pseudo-autosomal regions (PARs). PARs are regions of sex chromosomes of different types that can recombine (Otto et al., 2011); in humans, for example, there are two PARs, one at each end of the X and Y, that allow the X and Y to recombine with each other within that region (Helena Mangs & Morris, 2007). It is possible to incorporate PARs into a SLiM model with scripting (almost anything can be done with scripting!), but it is complex; an example is given in the SLiM manual. In the future, SLiM might be extended to support PARs intrinsically.

As discussed in the Introduction, in previous SLiM versions it was possible to model multiple diploid autosomes using single-position points in a recombination rate map with a rate of `0.5` so that the loci to the left and right of that point assort independently. This is still possible in SLiM 5, and the resulting statistics of inheritance are equivalent, so the question arises: when should this old technique be used, now that SLiM 5 supports multiple chromosomes? One scenario is when a large number of unlinked loci are modeled, each of which is perhaps just a single base position in length, as is common in theoretical models. Chromosomes are fairly heavyweight objects, in both runtime and memory usage, and so modeling a set of unlinked single-base-position loci as separate chromosomes will make SLiM do a great deal of extra work; and in any case, one might want more unlinked loci than SLiM 5’s limit of 256 chromosomes. The other common scenario where the `r=0.5` trick can be more performant is when the mutation rate is quite low, such that chromosomes are often partially or completely empty; in this case, SLiM’s internal `MutationRun` data structure, which optimizes for shared haplotype structure (as discussed further in SLiM’s manual), can then actually span more than one chromosome, taking advantage of the fact that the haplotype structure extends very broadly due to the lack of mutations in the model. There may be other cases where an `r=0.5` model outperforms a multi-chromosome model – although for the vast majority of the models in our performance test suite, SLiM 5 is faster than SLiM 4.x. Performance will also depend upon hardware and operating system details; in general, if performance is a key concern, one should test different approaches with a full-scale model to see empirically which approach is more performant.

An alternative to a multi-chromosome (or whole-genome) model is to separately model each chromosome as an independent run of SLiM (Rodrigues, Kern, & Ralph, 2024). This can have the advantage of utilizing multiple processing cores more completely in some cases, since runs for various chromosomes can be assigned to different cores. If the number of simulations to be run is large, however, this advantage would typically disappear, since each core could be occupied by a different run of the multi-chromosome model anyway. The disadvantage of this approach is in a loss of realism, since the simulations for different chromosomes follow different, independent genealogies, but linkage can extend across chromosomes. In some cases this might lead to extreme divergence between the runs for different chromosomes, such that aspects of their evolutionary dynamics such as population size and structure no longer match. It would be possible to avoid such issues by doing an initial run of the model with no genetics, recording the complete pedigree of the run and then exactly following that recorded pedigree in each per-chromosome run; however, this would be considerable additional work, and would usually only work for neutral models (for non-neutral models, the initial pedigree-generating run would generally need to include the multi-chromosome genetics, defeating the purpose).

In addition to developing new features for SLiM, we view documentation, education, and outreach as part of our task. We encourage new SLiM users who want to learn more about this software to take the SLiM Workshop; it is given occasionally in person, but there are also recorded lectures and PDF worksheets available online (https://messerlab.org/slim/#Workshops). The SLiM manual can also be a good way to learn; much of its length is in the form of more than 200 example “recipes” that show how to model many common scenarios, with explanation and discussion. The `slim-discuss` mailing list (https://groups.google.com/g/slim-discuss) is a great place to ask questions that go beyond what is provided in the manual.

Biology has seemingly endless complexities; supporting the modeling of full genomes is an important step towards allowing simulation-based study of a broader range of that complexity. We intend to continue maintaining SLiM and extending it with new features in the future, and we are very grateful for the support of the SLiM community in these ongoing efforts.

## Acknowledgements

Thanks to (in alphabetical order) Groves Dixon, Arielle Fogel, Gregor Gorjanc, Ben Jeffery, Jerome Kelleher, Anna Langmüller, Mikhail Matz, Manisha Munasinghe, Sohini Ramachandran, Elissa Sorojsrisom, and Melissa Wilson for helpful discussions, comments, and feedback, and in particular, to Tymoteusz Pieszko for coining the term “haplosome”. This work was supported by NIH awards R01HG012473 and R35GM152242.

## Data and Code Accessibility

SLiM 5 is open-source on GitHub at https://github.com/MesserLab/SLiM, and its home page is located at https://messerlab.org/slim/. Model code and data used to generate the figures is available at https://github.com/MesserLab/SLiM5Paper.

